# Path integration changes as a cognitive marker for vascular cognitive impairment? – a pilot study

**DOI:** 10.1101/2020.02.15.950675

**Authors:** Ellen Lowry, Vaisakh Puthusseryppady, Gillian Coughlan, Stephen Jeffs, Michael Hornberger

## Abstract

Path integration spatial navigation processes are emerging as promising cognitive markers for prodromal and clinical Alzheimer’s disease (AD). However, such path integration changes have been little explored in Vascular Cognitive Impairment (VCI), despite neurovascular change being a major contributing factor to dementia and potentially AD. In particular, the sensitivity and specificity of path integration impairments in VCI compared to AD is unclear. In the current pilot study, we explore path integration performance in AD and VCI patient groups and hypothesise that i) medial parietal mediated egocentric processes will be more affected in VCI and ii) medial temporal mediated allocentric processes will be more affected in AD. This retrospective cross-sectional study included early stage VCI patients (n=9), AD patients (n=10) and healthy age-matched controls (n=20). All participants underwent extensive neuropsychological testing, as well as spatial navigation testing. The spatial navigation tests included the virtual reality ‘Supermarket’ task assessing egocentric (body-based) and allocentric (map-based) navigation as well as the ‘Clock Orientation’ test assessing egocentric and path integration processes. Results showed that egocentric path integration processes are only impaired in VCI, potentially distinguishing it from AD. However, in contrast to our prediction, allocentric path integration was similarly impaired for VCI and AD. These preliminary findings suggest limited specificity of allocentric path integration deficits between VCI and AD. By contrast, egocentric path integration deficits emerge as more specific to VCI, potentially allowing for more specific diagnostic and treatment outcome measures for vascular impairment in dementia.

## INTRODUCTION

Vascular cognitive impairment (VCI) is the second most prevalent cause of cognitive decline after Alzheimer’s disease (AD) and is thought to account for ~20% of all dementias (Goodman et al., 2017; van der Flier et al., 2018). Although, individuals with mixed (AD and VCI) pathology are estimated to account for up to 70% of all dementia cases (Toledo et al., 2013). Despite the high prevalence of vascular impairment, its cognitive correlates are still being explored. Clinically, VCI is considered to involve a decline in executive function and higher order cognition such as information processing, planning, set-shifting and working memory (Hachinski et al., 2006; Sachdev et al., 2014). These changes are mostly attributed to micro and macro infarcts in subcortical and cortical regions, as well as their connecting white matter tracts (Beason-Held et al., 2012; van der Flier et al., 2018), in particular affecting fronto-parietal networks. Nevertheless, attributing such executive changes to VCI specifically has remained challenging, as executive function can also present as part of AD or related pathophysiology (Girard et al., 2013; Guarino et al., 2018; Neufang et al., 2011). However, the recent development of novel spatial navigation cognitive markers for AD show promise in being more specific to underlying disease pathophysiology (Coughlan et al., 2018a) and may help to identify cognitive decline specific to VCI. A clear distinction between VCI and AD is critical as with appropriate intervention VCI can be slowed or halted whereas AD has a fixed and terminal prognosis.

Spatial navigation is a fundamental cognitive skill that requires the integration of egocentric (body-based) and allocentric (map-based) frames of orientation. Both frames are required for everyday navigation with egocentric and allocentric processes shifting as a function of navigational demands (McNaughton et al., 2006). Path integration is integral to spatial navigation as it allows an individual to keep track of and return to their starting location on the basis of visual, self-motion, vestibular and proprioceptive feedback which represent current position and heading direction in references to a permanent location (Etienne and Jeffery, 2004: Knierim, Neunuebel and Deshmukh, 2014; McNaughton et al., 2006). This process involves translating distance travelled with changes in direction of movement either relative to our allocentric or egocentric orientation (Burgess, 2006). Multisensory (visual, self-motion, vestibular and proprioceptive) feedback combine egocentric and allocentric frames of reference, allowing path integration to continuously update this information, allowing one to keep track of one’s position in space (Coughlan et al., 2018a; Rieser, 1989).

Egocentric orientation relies more on the prefrontal and parietal cortex to localise the position of objects relative to the body (Arnold, Burles, bray, Levy and Giuseppe, 2014; Goodale & Milner, 1992), the precuneus then uses these location cues to form the basis of an egocentric representation of the surrounding space, integrating self-motion cues with the egocentric reference frame (Woblers and Weiner, 2014). While, allocentric orientation is reliant on the formation of maps using place, grid and boundary vector cells situated mainly in the medial temporal lobe (Coughlan et al., 2018a; Lester et al., 2017). The integration of egocentric and allocentric frames occurs in the retrosplenial cortex (RSC), which is a critical interface between the medial temporal and medial parietal regions (Alexander & Nitz, 2015). Dorsal-medial regions of the RSC are thought to be implicated in orientating and recalling unseen locations from a current position in space, whilst ventro-lateral portions were more linked to updating and integrating scene information (Burles, Slone and Giuseppe, 2017).

Tasks that tap into path integration therefore provide a promising ecological, cognitive framework to detect medial temporal and medial parietal pathophysiology. Not surprisingly, path integration has been already explored in AD (Morganti et al, 2013; Ritchie, 2018; Serino et al., 2014; Vlcek & Laczo, 2014) and the advent of VR based testing has allowed such tests to be clinically available (Morganti et al., 2013; Parizkova et al., 2018; Plancher et al., 2012). We have developed previously such a test, the Virtual Supermarket task, which is now used across many large cohorts and drug trials as it can reliably detect path integration differences in preclinical and clinical dementia populations (Tu et al., 2017; Tu et al., 2015). The VR task reliably measures spatial processes of: i) egocentric self-reference navigation; ii) allocentric map-based navigation and iii) heading direction. For example, we have previously shown that the test allows distinction of behavioural variant fronto-temporal dementia (bvFTD) from AD, with AD showing particularly problems in switching between egocentric and allocentric frames during path integration (Tu et al., 2017). Importantly, these switching problems in AD were associated with grey matter atrophy in the RSC (Tu et al., 2015).

In contrast to the exciting findings in AD, less is known about path integration in VCI, despite path integration potentially allowing as well to tap into parietal deficits in VCI (Haight et al., 2015; Maguire, 1998; Papma et al., 2012; Wolbers et al., 2004). A previous case study by our group explored path integration in a 65 year old male with VCI. The findings showed that the vascular patient had normal performance on allocentric orientation but a clear and isolated deficit in egocentric and heading direction sub-components of the path integration tasks (Coughlan et al., 2018b). These findings are consistent with fronto-parietal network disruptions typically seen in vascular dementia patients (Beason-Held et al., 2012; Sachdev et al., 2014; van der Flier et al., 2018) and may suggest medial parietal changes imped the egocentric frame of reference and subsequent path integration.

The current study leads on from this case study by exploring path integration in a group of VCI patients, and importantly comparing them against a group of AD patients and controls. Navigation will be tested using the Virtual Supermarket task where participants move through the virtual environment to a series of locations and are tested on their egocentric, allocentric and heading direction response. We hypothesise that i) medial parietal mediated egocentric processes will be more affected in VCI; ii) medial temporal mediated allocentric processes will be more affected in AD.

## MATERIALS AND METHODS

### Participants

Nine vascular cognitive impairment and 10 Alzheimer’s disease patients along with 20 healthy controls were recruited to participate in a research study at the University of East Anglia as part of the wider The Dementia Research and Care Clinic (TRACC) study. The study was approved by the Faculty of Medicine and Health Sciences Ethics Committee at the University of East Anglia (reference 16/LO/1366) and written informed consent was obtained from all participants. Clinical diagnosis (VCI or AD) was classified by a consultant at the Norfolk and Suffolk Foundation Trust by interviewing the patient, examining neuropsychological assessment scores, structural clinical MRI scans and the patient’s medical history. Disease duration was reported by the person’s study partner (a spouse or relative). Participants had no history of psychiatric or neurological disease, substance dependence disorder or traumatic brain injury and had normal or corrected-to-normal vision. All participants underwent neuropsychological screening, including cognitive screening, episodic memory and spatial memory tasks, Addenbrooke’s cognitive examination (ACE-III), Rey–Osterrieth Complex Figure Test (RCFT) copy and with 3-min delayed recall, Cube Analysis, Dot Counting and Position Discrimination from the Visual Object and Space Perception Battery (VOSP).

### Virtual Supermarket Task

The Virtual Supermarket Task has been developed by our group previously and used in symptomatic mild cognitive impairment (MCI), AD, frontotemporal dementia (FTD) and VCI patients (Coughlan et al., 2018b; Tu et al., 2017; Tu et al., 2015). The VR task is an ecological test of spatial navigation abilities designed to simulate navigating through a real-world supermarket. An iPad 9.7 (Apple Inc.,) was used to show participants 20-40 second video clips of a moving shopping trolley in the virtual supermarket (Figure 1A-C). Videos were presented in a first-person perspective and participants are provided with optic flow cues from the moving shopping trolley and changing scenery as it followed different routes to reach a different end point in each trial. The task avoids the use of landmarks or salient features within the environment and limits the demand on episodic memory, reflecting similar tasks in the literature (see, Cushman, Stein and Duffy, 2008; Woblers, Weiner, Mallot and Büchel, 2017; Serino, Morganti, Di Stefano and Riva, 2015). The test taps into path integration processes via three core spatial processes: i) egocentric self-reference navigation; ii) allocentric map-based navigation and iii) heading direction. Once the video clip stops, participants indicate in real-life the direction of their starting point (egocentric orientation; Figure 1D). In a second step, participants indicate their finishing location on a birds-eye view map of the supermarket (allocentric orientation; Figure 1E), performance is calculated using the distance error (mm) between this and the coordinates of the actual finishing location. This map-based component provides an assessment of geocentric encoding of the virtual environment. The participant then indicates their heading direction at the finishing point, which determines the ability to which heading direction was encoded and updated throughout the task. The tasks consists of 14 trials and takes approximately 10 minutes to complete.

### Clock Orientation test

The Clock Orientation test has also been developed by our lab (Coughlan et al., 2018b) as a bedside clinical test for egocentric orientation. It requires participants to imagine they are standing in the centre of a large clock, facing a particular number, e.g., the number 3.

Participants are then asked “which number is directly behind you?” (Answer: number 9). Next participants are asked to point, in real-life, to the positions of different numbers on the clock face in relation to the number that they are currently facing. For example, “You are facing number 12, can you point to the number 3?” (Answer: pointing right). The questions increase in complexity across the test and require medial parietal mediated mental imagery, rotation and egocentric processes, with no episodic memory demand. The test consists of 12 trials and takes 5-10 minutes to complete.

### Procedure

Participants completed a battery of neuropsychological assessments at their home (see Table 1 for list of tasks). In a second session held at the Norfolk and Suffolk Foundation Trust, participants undertook cognitive experimental tests (including the virtual Supermarket task and Clock Orientation test) and completed a clinical interview with the Chief Investigator of the study.

### Statistical Analysis

Statistical analysis was performed using IBM SPSS (Version 25). Chi square and two tailed one-way univariate analysis of variance (ANOVA) were used to test the significance of any demographic or neuropsychological differences between the clinical groups. When quantifying group differences, partial eta squared (*n_p_^2^*) was used as a measure of effect size. The Supermarket task has 3 measures-specifically egocentric response, allocentric response and heading direction. Each outcome measure was individually entered into a one-way analysis of covariance (ANCOVA) with group as the independent variable and age and sex as covariates. The Clock Orientation test was also analysed using a one-way ANCOVA with group as the independent variable and age and sex as covariates. Post-hoc pairwise comparisons were conducted using Bonferroni adjustment for multiple comparisons. Sensitivity and specificity of the egocentric supermarket task and clock orientation test performance in VCI and AD were compared using logistic regression and ROC curve analysis. A Z-score of AD performance was computed for 7 missing values for one AD patient in the Virtual Supermarket test.

**Figure 1.**
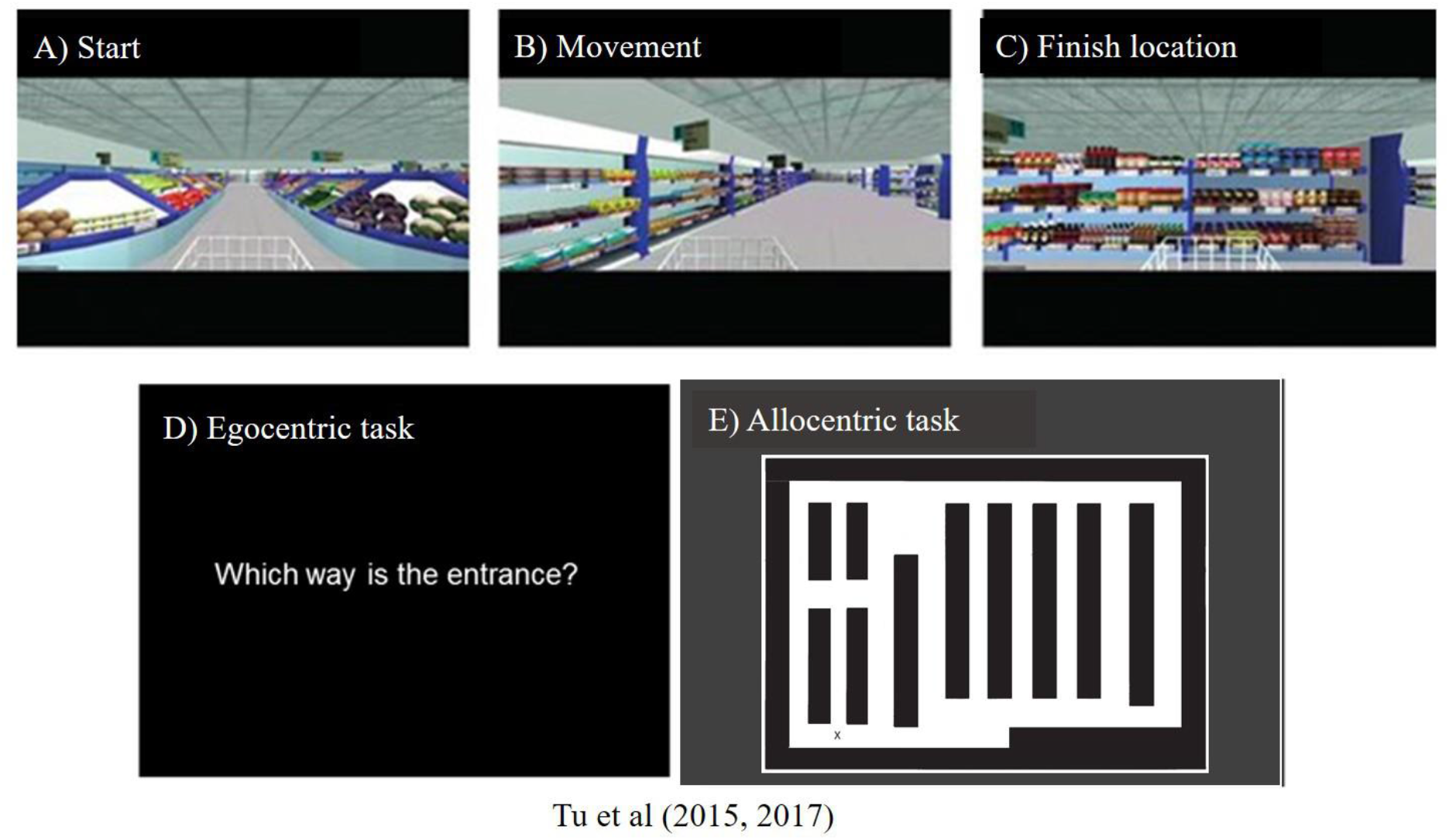
Screenshots from the supermarket task, showing A) starting viewpoint, B) movement during example video clip, C) end location of an example video clip, D) onscreen instructions prompting participant to indicate direction of their starting point, E) the supermarket map participants use to indicate their finishing location and their heading direction when the video clip ends.

## RESULTS

### Demographics and Neuropsychology

Participant groups were well matched and no significant differences in demographic measures were observed between the VCI, AD and control groups (all p-values > .1). ANOVA of participant groups showed both VCI and AD patients performed significantly lower on a general cognitive screening test (ACE-III) and the memory recall domain of RCFT compared to controls (all p-values < .01). Results showed no significant neuropsychological differences between the VCI and AD patients for the ACE-III, RCFT recall condition, VOSP dot counting and cube analysis sub-sets (all p-values > 0.1. However, VCI patients were significantly more impaired than AD patients in the RCFT copy condition, FCSRT free recall condition and the VOSP position discrimination (all p-values < .1) (see table 1).

**Table 1.** Demographic characteristics and Neuropsychological Performance.

|  | VCI | AD | Control |  |
| --- | --- | --- | --- | --- |
|  | Mean (SD) | Mean (SD) | Mean (SD) | Sig post-hoc VCI vs. AD comparisons |
| *n* | 9 | 10 | 20 |  |
| Sex (F/M) | 3/6 | 2/8 | 9/11 | *ns* |
| Age | 70.22 (4.57) | 69.91 (7.7) | 69.6 (6.45) | *ns* |
| Disease duration | 3.13 (2.64) | 2.81 (2.21) | n/a | *ns* |
| **General cognition** |  |  |  |  |
| Total ACE-III | 69.44 (12.9) | 72.1 (22.41) | 95.1 (3.13) | *ns* |
| ACE: Attention | 13.5 (.72) | 15.75 (.72) | 17.6 (.45) | *ns* |
| ACE: Memory | 13.5 (1.73) | 17.13 (1.17) | 24.3 (.74) | *ns* |
| ACE: Fluency | 7.13 (.59) | 8.12 (.59) | 11.7 (.37) | *ns* |
| ACE: Language | 21.77 (2.44) | 22.33 (3.04) | 25.6 (.61) | *ns* |
| ACE: Visuospatial | 11.5 (1.19) | 16.67 (1.12) | 15.8 (.75) | * |
| **Visuospatial ability** |  |  |  |  |
| RCFT: Copy | 22.1 (7.17) | 28.4 (8.92) | 32.72 (3.23) | * |
| RCFT: Recall | 7 (5.65) | 11.8 (8.12) | 17.55 (5.43) | *ns* |
| Dot Counting | 9.5 (0.71) | 9.8 (0.42) | 10 (0) | *ns* |
| Position Discrim | 18.87 (1.27) | 19.7 (0.67) | 19.85 (0.37) | * |
| Cube Analysis | 8.11 (2.62) | 8.7 (1.88) | 9.8 (0.52) | *ns* |
| **Memory ability** |  |  |  |  |
| Total FCSRT | 29.21 (2.84) | 42.91 (2.63) | 47.92 (2.01) | ** |
| FCSRT: Free recall | 8.83 (7.94) | 17.14 (8.83) | 26.83 (4.17) | *ns* |
| FCSRT:Cued recall | 25.7 (4.94) | 20.5 (7.2) | 23.35 (4.87) | *ns* |
| **Supermarket test** |  |  |  |  |
| Egocentric | 3.44 (3.24) | 9.4 (2.27) | 8.1 (3.7) | ** |
| Allocentric | 69.1 (38.11) | 48.41 (12.17) | 30.2 (14.13) | *ns* |
| Head direction | 4.8 (1.33) | 5 (3.41) | 7.1 (0.9) | *ns* |
| **Clock test** | 5.43 (0.81) | 10.1 (1.2) | 10.1 (0.51) | *** |

\* Significant group differences between VCI and AD patients. \* $p < .1$ , \*\* $p < .01$ , \*\*\* $p < .001$ . ACE-III= Addenbrooke's cognitive examination. RCFT: Copy= Rey-Osterrieth Complex Figure Task, copy condition. RCFT: Recall= Rey-Osterrieth Complex Figure Task, recall 3 minutes after copy. Dot Counting, Position Discrimination and Cube Analysis= sub-sets from Visual Object and Space Perception Battery (VOSP). FCSRT: free recall= Free and Cued Selective Reminding, free recall Test condition, FCSRT: free recall= Cued and Cued Selective Reminding Test, cued condition.

### Virtual Supermarket Task

An ANCOVA with age and gender as covariates revealed a significant differences between egocentric responses on the supermarket test, *F(2, 34) = 8.14, p < .001, n_p_^2^ = .32*. Post-hoc comparisons revealed significantly greater egocentric impairment in VCI (M= 3.5, SD= 3.24) compared to AD (M= 10.01, SE= 1.11), *p* < .002, 95% CI [-10, −2.1] and control groups (M=8.1, SD= 3.7),*p* < .009, 95% CI [-7.95, −1.1]. No other significant group differences were observed (*p* > .1) (see figure 2A).

Allocentric responses showed a significance difference between groups, controlled for age and gender *F(2,34) = 10.1, p <.001, n_p_^2^ = .37*. Post-hoc comparisons showed significantly greater impairments in VCI patients (M = 68.33, SD= 38.1) compared to controls (M= 30.85, SD= 14.13), *p* < .001, 95% CI [16.02, 61.1] but impairments did not reach statistical significance in AD patients (M= 50.1, SD= 7), *p* = 0.09, 95% CI [-41.11, 2.1] compared to controls. However, there were no significant groups differences between VCI and AD (p>.1) (see figure 2B).

Heading direction (correct judgement of facing direction after travel period) did not reveal significant group differences when controlling for age and gender *F(2, 34) = 1.11, p > .1, n_p_^2^ = .06* (see figure 2C).

### Clock Orientation Test

An ANCOVA with age and gender as covariates revealed a significant difference between egocentric responses on the Clock Orientation task *F(2, 34) = 13.4, p < .001, n_p_^2^ = .44.* Post-hoc comparisons showed significantly greater egocentric deficits in VCI patients (M= 5.42, SD= 3.16) compared to AD (M= 10.1, SD= 1.21), p < .001, 95% CI [-7.2, −2] and control groups (M= 9.65, SD= 2.06), p < .001, 95% CI [-6.56, −7.1]. No other significant group differences were observed (p > .1) (see figure 2D).

**Figure 2.**
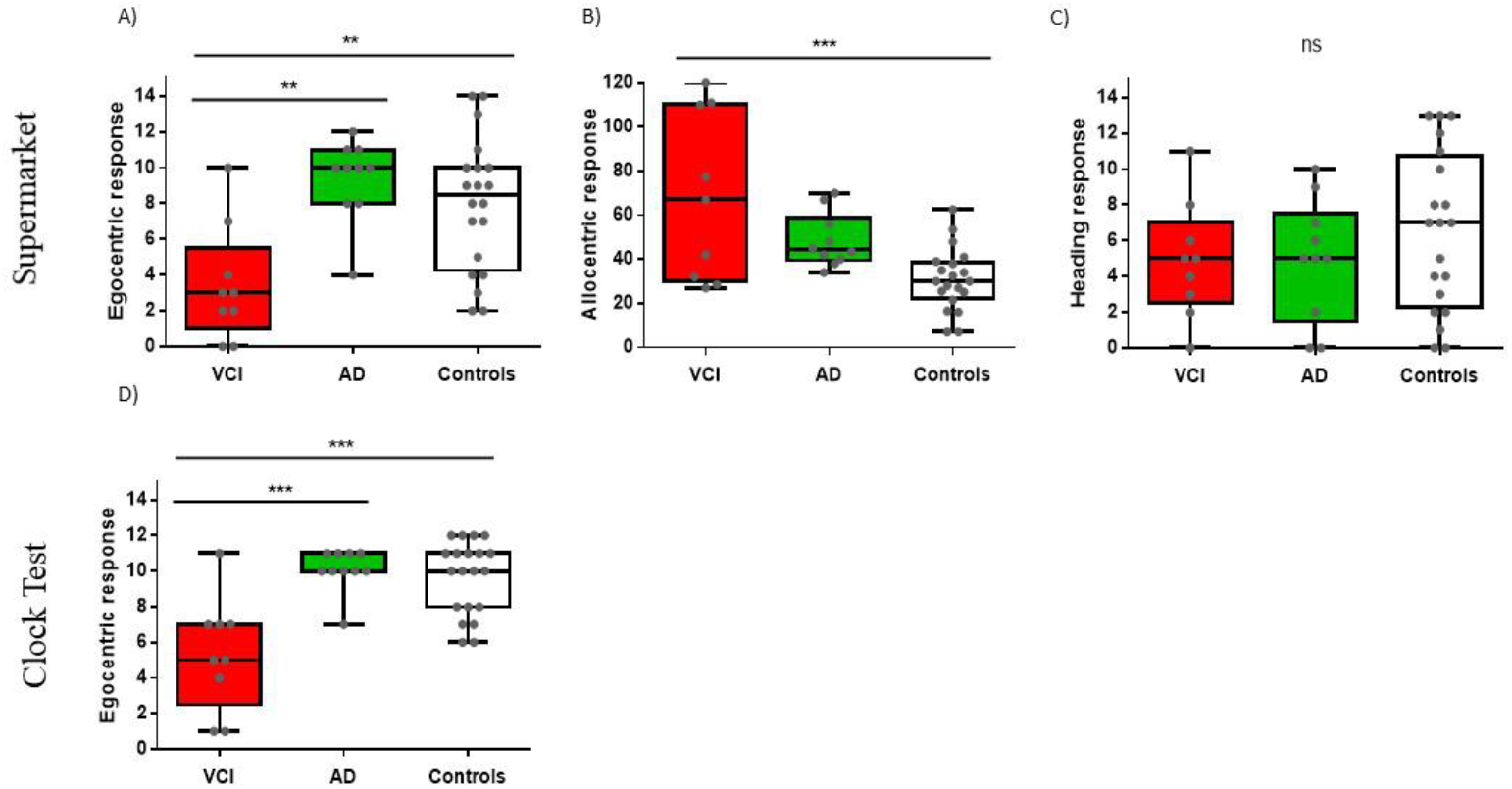
Spatial orientation performance between VCI, AD and Controls. \*\**p*<.0l, \*\*\**p*<.00l. Supermarket task displays Egocentric response (correct), Allocentric response (error in mm) and Heading response (correct). Clock Orientation test displays Egocentric response (correct).

### Sensitivity and Specificity

Sensitivity and specificity of egocentric supermarket and clock test performance in VCI and AD were explored using logistic regression and ROC curves. Logistic regression indicated that the regression model based on egocentric scores of Supermarket and Clock Orientation predictors was statistically significant, X^2^(2) = 16.36,*p* < .001. The model explained 77% (Nagelkerke R^*2*^) of variance in VCI and AD patients and correctly classified 84% of patients (7 out of 9 VCI; 9 out of 10 AD) into their respective cohorts. ROC curves were computed for the supermarket and clock test predictors in discerning VCI from AD patients. Similarly, Area Under the Curve (AUC) values indicated that egocentric orientation in the Supermarket (AUC = .8, SE = .12; 95% CI [.56, 1]) and Clock test (AUC= .91, SE = .06, 95% CI [.8, 1]) had strong diagnostic accuracy in distinguishing VCI from AD patients.

**Figure 3.**
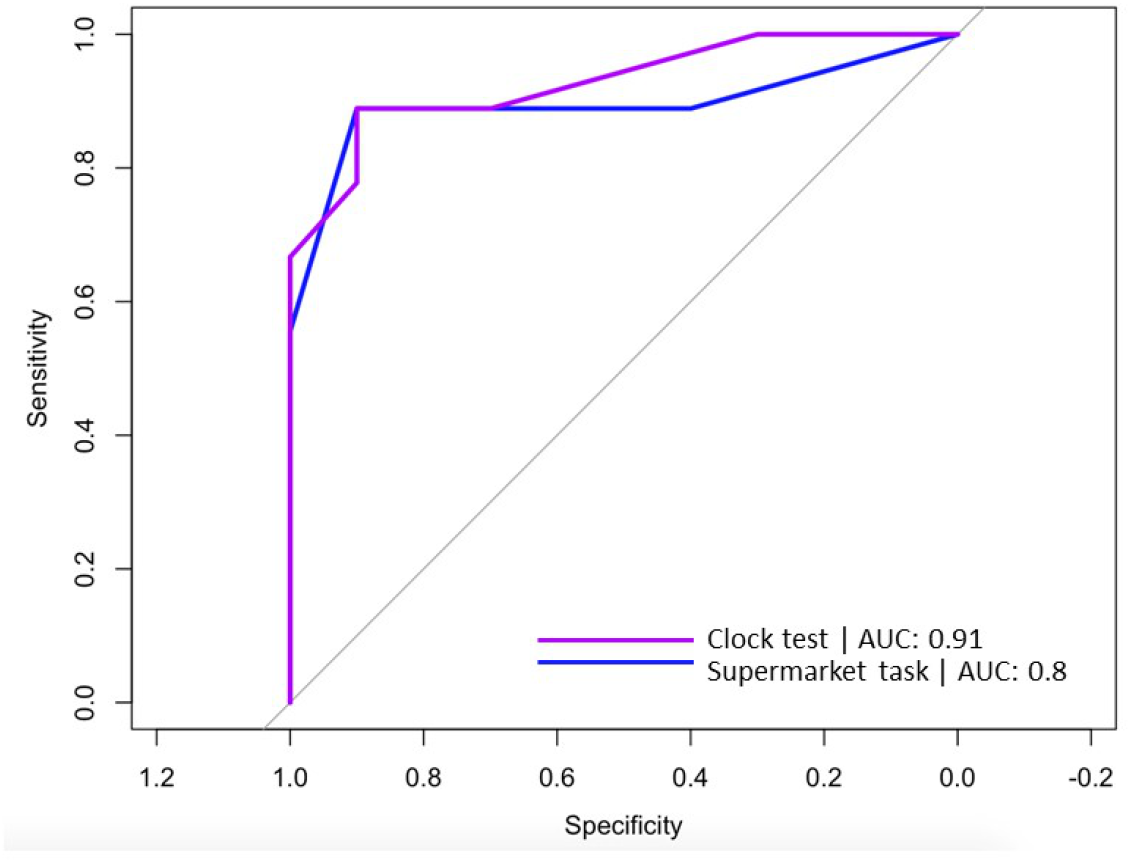
ROC curves for Supermarket task (blue line) and Clock test (purple line) predicting correct diagnosis (VCI or AD).

## DISCUSSION

Overall, our results indicate that medial parietal mediated egocentric path integration processes are a sensitive and specific cognitive marker selective for VCI. By contrast, allocentric orientation deficits were less sensitive, and not specific to distinguish between the underlying pathologies.

In more detail, the egocentric path integration measures of the Virtual Supermarket task and Clock Orientation test successfully detect vascular changes in patient populations. More importantly, the measures allowed to reliably distinguish vascular from AD pathophysiology in the patient populations. Notably, egocentric orientation was impaired in VCI, but relatively intact in AD patient groups when controlling for age and gender. This supports findings from our vascular patient case study (Coughlan et al., 2018b) and suggests egocentric impairments indicate a more medial parietal focused change (Weniger et al., 2009) in VCI. Furthermore, the AD patient’s egocentric ability remained intact which supports suggestions that MCI and earlier stage AD groups show an undisturbed egocentric orientation (Coughlan et al., 2019). It would be interesting to explore whether more moderate to advanced AD patients might show problems using both allocentric and egocentric orientation, as it is known that medial parietal structures might be affected only later in the disease course (Braak & Del Tredici, 2015).

The egocentric demands in the virtual Supermarket requires the individual to form an accurate representation of the starting point by integrating virtual self-motion with heading direction to reach their end destination. Path integration plays an important role in updating spatial orientation during self-motion but this process is accumulative, therefore can be liable to directional errors with respect to the original starting position (McNaughton et al., 2006), which may be responsible for problems observed across both egocentric tasks. The Clock Orientation test also demands path integration to configure the position of numbers on a clock face relative to the individual’s current position. Both tasks rely on accessing scene construction, mental rotation and imagery translated from an egocentric orientation. At the neural level, translation of these egocentric processes depend mainly on medial parietal cortex (Coughlan et al., 2018a; Galati et al., 2000; Goodale & Milner, 1992; Zaehle et al., 2007) as well as prefrontal cortex (Bird et al., 2012; Spiers, 2008; Spiers & Barry, 2015), indicating potential disruptions in fronto-parietal structures typically seen in vascular patients (Beason-Held et al., 2012; Heiss et al., 2016; van der Flier et al., 2018; Vipin et al., 2018).

Medial parietal mediated egocentric deficits appear to characterise VCI patients. This is consistent with emerging evidence suggesting the earliest signs of dysfunction appear in medial frontal and anterior cingulate regions in at VCI-risk individuals (Haight et al., 2015; Papma et al., 2012), which is accompanied by a more typical vascular profile of reduced integrity of white matter in the bilateral superior longitudinal fasciculus (Beason-Held et al., 2012). Since egocentric orientation does not deteriorate in healthy aging and early stage AD, compared to medial temporal based cognitive functions (for review, see Colombo et al., 2017) it emerges as a potential powerful cognitive marker to identify early vascular-related pathology. Given the prevalence of vascular related dementia it is surprising that investigation to isolate cognitive deficits unique to this pathology is so sparse. However, based on our findings, it appears that egocentric orientation may be a useful diagnostic tool to discriminate VCI from other neurodegenerative conditions.

Our study suggests allocentric orientation deficits were not statistically present in AD, only VCI showed significant impairments compared to healthy controls. This does not support our prediction that allocentric deficits would be more profound in AD. The literature suggests allocentric deficits are more prominent in preclinical AD (Coughlan et al., 2019) with a loss in selectivity as the disease stage progresses and deficits become more widespread (Braak & Del Tredici, 2015). Yet, in the early stage AD patients in our study results did not reach significance. One potential explanation for the results observed may be provided by the large range in allocentric scores across the VCI group. VCI is a highly heterogeneous disordered in terms of disease pathology and subsequent cognitive impairments which may account for this variation, compared AD pathology and symptoms are more uniform. Indeed, as evident from Figure 2, it is clear that AD patients perform differently from controls but this did not reach statistical significance. VCI patients revealed both egocentric and allocentric orientation problems which is likely to represent a disruption to translational and integration processes where both frames are combined to produce effective navigation. This view also explains the reduced visuospatial performance exhibited by the VCI patients during neuropsychological testing across RCFT copy and position discrimination tasks. It is also important to consider the domain of memory when interpreting our findings. Results from the FCSRT suggest VCI patients had significantly worse memory than the AD and control groups, sub-score results indicate this is driven by reduced performance during free recall. This is likely due to the retrieval demands on pre-frontal and parietal structures (Staresina and Davachi, 2006) which are typically disrupted in VCI. However, when cued VCI patients outperform AD patients. This finding is consistent with evidence that suggests providing a cue has little bearing on improved memory recall in AD (Sarazin et al., 2007; Wagner et al., 2012). This may be relevant to the poor allocentric results observed for VCI patients, as reduced retrieval mechanisms may have disrupted their task performance opposed to pure allocentric (medial temporal) mapping problems, which we would expect to see in the AD patients.

Despite these exciting findings, our study is not without limitations. First and foremost, the sample sizes for the groups were small and therefore replication in larger patient cohorts would be important. Further, clinical characterisation of VCI subtypes (Skrobot et al., 2017) would help to better classify vascular pathology and determine accompanying cognitive symptoms, this may also help inform the variation of results seen in allocentric performance for the VCI patients. Finally, we did not have neuroimaging biomarker confirmation of vascular or AD pathophysiology. Confirmation of vascular lesions and their locations, as well as AD specific biomarkers would be important in the future to corroborate our cognitive findings.

Nevertheless, to our knowledge this in the first study to isolate a selective navigational deficit in VCI. This showcases the important role of virtual navigation and spatial tests in the future development of sensitive and specific diagnostic tests for VCI. Further investigation into the cognitive symptoms selective to VCI as well as longitudinal cohort studies in at VCI-risk individuals is critical to identify the emergence of the disease and intervene with therapeutic strategies as early as possible.

In conclusion, our findings show a distinct egocentric orientation deficit that is specific for VCI relative to AD. This is critical given the lack of specificity in current diagnostic tests and the indistinct diagnostic criteria for cognitive symptoms in VCI. In turn, this will inform diagnostic work-ups and aid personalised treatment pathways to treat underlying vascular changes in patients.

## Abbreviations

VCI: vascular cognitive impairment,
AD: Alzheimer’s disease

## AUTHOR CONTRIBUTION

EL and MH contributed to the conception and design of the study, statistical analysis and the intellectual contribution to the writing of the manuscript. VP, GC and SJ contributed to the data collection and intellectual contribution to the manuscript.

## ACKNOWLEDGEMENTS

We would like to thank the Norfolk and Suffolk Foundation Trust and the patients and families involved in the TRACC study.

